# Automatic identification of informative regions with epigenomic changes associated to hematopoiesis

**DOI:** 10.1101/082917

**Authors:** Enrique Carrillo-de-Santa-Pau, David Juan, Vera Pancaldi, Felipe Were, Ignacio Martin-Subero, Daniel Rico, Alfonso Valencia, on behalf of The BLUEPRINT Consortium

**Author notes:** Co-senior and corresponding authors. Co-first authors (Enrique Carrillo-de-Santa-Pau, David Juan).

## Abstract

Hematopoiesis is one of the best characterized biological systems but the connection between chromatin changes and lineage differentiation is not yet well understood. We have developed a bioinformatic workflow to generate a chromatin space that allows to classify forty-two human healthy blood epigenomes from the BLUEPRINT, NIH ROADMAP and ENCODE consortia by their cell type. This approach let us to distinguish different cells types based on their epigenomic profiles, thus recapitulating important aspects of human hematopoiesis. The analysis of the orthogonal dimension of the chromatin space identify 32,662 chromatin determinant regions (CDRs), genomic regions with different epigenetic characteristics between the cell types. Functional analysis revealed that these regions are linked with cell identities. The inclusion of leukemia epigenomes in the healthy hematological chromatin sample space gives us insights on the healthy cell types that are more epigenetically similar to the disease samples. Further analysis of tumoral epigenetic alterations in hematopoietic CDRs points to sets of genes that are tightly regulated in leukemic transformations and commonly mutated in other tumors. Our method provides an analytical approach to study the relationship between epigenomic changes and cell lineage differentiation. Method availability: https://github.com/david-juan/ChromDet

## Introduction

Hematopoiesis is one of the most studied biological differentiation processes, in which different cell lineages arise from a common hematopoietic stem cell (HSC). This system can be seen as a hierarchical tree, where the more internal ‘nodes’ are the different lineage progenitors and the ‘leaves’ are the final mature cell types (1, 2). This hierarchical tree with many ‘nodes’ and ‘leaves’ provides the best model to study chromatin remodeling during cell lineage differentiation (3–5).

Chromatin remodeling is a dynamic process that modulates the chromatin architecture and is vital to ensure proper functioning of the cell and maintenance of its identity (6). The deregulation of chromatin remodeling factors often leads to diseases such as cancers (7) and neurodevelopmental disorders (8, 9). A main role in this re-organization of chromatin is played by post-translational modifications of histone tails, which can affect many biological processes such as gene transcription, DNA repair, replication and recombination (10), (11). Moreover, the cross-talk between different modifications affects the binding and function of other epigenetic elements, increasing the complexity of the chromatin remodeling process (12).

Despite great progress in our understanding of hematopoiesis during the last decades (13, 14), we are still far from fully uncovering the details of the epigenetic mechanisms controlling this process. It is now widely accepted that the cell phenotype is directly related to its epigenetic makeup and that chromatin changes during differentiation contribute to the determination of cell fate. However, a major challenge in the field is to identify exactly where the epigenetic changes causing phenotypic changes occur. Similarly to the problem of distinguishing driver and passenger mutations in cancer, we can think of driver and passenger chromatin changes during cellular differentiation. Chromatin drivers of cellular differentiation would correspond to the subset of regions whose change is required to perform the different differentiation steps. As consequence, these regions must reflect one or more changes among cell types, while being fixed in any specific cell type. We therefore advocate the need to develop strategies identifying these key chromatin regions and their epigenetic changes that drive differentiation and determine cell fate. For this purpose, we take advantage of the large and comprehensive epigenomics datasets produced by the partners of the International Human Epigenome Consortium (IHEC; http://ihec-epigenomes.org/).

Here, we propose an approach to identify the key chromatin regions that undergo chromatin changes associated to cell differentiation during multiple differentiation steps in hematopoiesis. We define chromatin states based on the combinatorial patterns of 6 histone modifications in 42 human samples covering the myeloid and lymphoid differentiation lineages from hematopoietic stem cells (HSCs). This framework establishes highly informative low-dimensional spaces based on a Multiple Correspondence Analysis (MCA; (15)) of the profiles of histone modification combinations (chromatin states). Our integrative analysis of chromatin states in these samples recapitulates the human hematopoietic lineage differentiation tree from an epigenetic perspective. Moreover, our approach identifies 32,662 chromatin determinant regions (CDRs) in which chromatin changes are associated with the various differentiation steps the cells go through, possibly influencing their final lineage identities. The combination of chromatin states in these CDRs constitutes an epigenomic fingerprint that characterizes the different hematopoietic cell types. The method is available at https://github.com/david-juan/ChromDet

## Material and Methods

### ChIP-Seq Data Processing

We retrieved data for 430 chromatin immunoprecipitation sequencing (ChIP-Seq) experiments from BLUEPRINT, ENCODE and NIH ROADMAP. We downloaded the hg19/GRCh37 alignments for 2 CD4+ and 1 CD8+ lymphocytes, 5 mature neutrophils, 3 CD14+ monocytes, 4 macrophages and 7 CD38-B cell samples from BLUEPRINT; 11 CD4+ and 3 CD8+ lymphocytes, 1 CD14+ monocyte and 3 CD34+ hematopoietic stem cells samples from NIH ROADMAP and 2 CD14+ monocytes samples from ENCODE described in **S1 Table** and Fig. 2A. In addition, the analysis including diseases was based on data from 3 Acute Myeloid Leukemias (AML), 6 Chronic Lymphocytic Leukemias (CLL) and 3 Mantle Cell Lymphomas (MCL) from BLUEPRINT (see **S1 Table**). The BAM files were converted to BED format and duplicate reads were removed for all the experiments. We computed different quality control measures with phantompeakqualtools (v1.10.1; see (16)) including total number of reads, normalized strand cross-correlation coefficient (NSC) and quality tag based on thresholded relative strand cross-correlation coefficient (RSC; see **S1 Table**). We flagged those histone experiments with less than 10^7^ reads and no replicates; NSC <1.05 and quality tag based on RSC <0. Then, following a similar strategy used previously by the NIH Epigenomics Roadmap (see (17)), we computed an overall quality rating per sample based on the six core histone modification quality experiments. We labelled samples as “very high quality” if none or only 1 histone mark experiment failed in 1 out of the 3 quality criteria; “high quality” if 2 or 3 histone experiments failed in 1 out of the 3 quality criteria; “medium quality” if more than 3 histone marks failed in 1 out of the 3 criteria or up to two broad histone modifications (H3K36me3; H3K9me9; H3K27me3) failed in 2 out of the 3 quality criteria; “low quality” if 3 or more histone experiments failed in 2 out of the 3 quality criteria or at least 1 histone experiment failed in the 3 quality criteria used. All the samples labelled as “low quality” were discarded and not included in our study. The overall quality criteria for those histone experiments included in the analysis is shown in **S1 Table**.

### Genome Segmentation

The input information used to segment the genome into different chromatin states was derived from 6 histone modifications (H3K4me3; H3K4me1; H3K27me3; H3K9me3; H3K27ac and H3K36me3). We used the ChromHmm software (v1.10;(18)) to define a 11 chromatin-states model (see **S1 Fig**) following the strategy proposed by the ChromHMM developers to set up the different parameters like number of states for training or posterior collapse (17, 19, 20). We evaluated the consistency and interpretability of chromatin states in models learnt with different numbers of chromatin states (5, 7, 9, 11, 13 and 20 states), quantified as the correlation of chromatin mark frequencies obtained for corresponding states across different models, as previously done by Ernst M., and Kellis M. (19). The results show that the 20 states model is recovered with correlations higher than 0.75 by the states trained in the model with 11 states, with little improvement in the 13 states model (**S2-S3 Tables**). The 11 states model captures all the biological-interpretable states that were consistently found in larger models.

Importantly, a manual curation of the chromatin states based on available additional information (gene structures, CpG islands, Lamin B1, etc) showed that the 11 states model retrieves all the main regulatory states (active promoter, bivalent promoter, enhancer, elongation, heterochromatin/low signal), without including any functionally unclassifiable chromatin state. Therefore, our approach of selecting 11 states to train the HMM is aimed at striking an equilibrium between a low enough number of combinations and the biological interpretability of the states included in the analysis, based on the ChromHMM emission probabilities correlation, prior knowledge regarding the function of these marks, and our previous experience (12). In summary, the 11 states model selected captures the biologically-interpretable states that were consistently found in larger models providing a suitable framework for our analysis.

We generated the model with the “healthy” samples excluding B cells (see **S1 Table** for details). The samples from B cells (naive and tonsil) and diseases (AML, CLL, MCL) were segmented with the model generated previously, as they were produced at the final stages of the BLUEPRINT project. Further, segmentations for each sample from the 11 states model were collapsed into 5 chromatin states summarizing similar states based on the emission probabilities, literature, biological knowledge, and genomic feature enrichments: heterochromatin/low signal (H), enhancer (E), transcription (T), active promoter (A) and repressed promoter (R; **S1A Fig**). Our a posteriori collapse into 5 chromatin states let us group dynamic states for a more robust representation of the epigenomic variability in cell types. In fact, differences in strength of enhancers, promoters or elongating regions can reflect more or less dynamic regions resulting in subtle differences between ChIP-seq experiments.

Therefore, for each sample, we have a vector of regions with their corresponding labels (chromatin states). In addition, we partitioned the genome into 200bp, preserving the associated chromatin state labels in order to have the same number of regions in all samples and make them comparable. For further analysis, consecutive 200bp intervals with the same labels pattern in all samples were merged, any change in one sample marking an interval transition.

### Sample clustering in the Chromatin Sample Space obtained by Multiple Correspondence Analysis (MCA)

In this work we propose to use a methodological protocol based on Multiple Correspondence Analysis (MCA; (15)), previously applied to multiple sequence alignments of proteins, for the automatic extraction of relevant signatures (21) and to gene expression profiles for sample classification (22). MCA can be considered as an equivalent to Principal Component Analysis (PCA) when working with qualitative data instead of continuous variables. MCA disentangles the sources of epigenomic variability among our samples into a set of principal components that form an orthogonal space which dimensions can be prioritized according to their corresponding eigenvalues. This MCA space can be reduced to a low dimensional one preserving most of the original information but filtering the main sources of noise. In brief, our protocol performs a MCA on a vectorial representation of multiple chromatin states sample vectors. It establishes the informative low dimensional space incorporating only those components with the highest eigenvalues, those explaining most of the total variance, where samples coordinates distribution is statistically different (P-value < 0.01, Wilcoxon test) between the tested component and the previously selected one, the one with the closer higher eigenvalue. In this work, we define the Chromatin Sample Space as the space formed by this set of highly informative components coming from the MCA on the vectors of the chromatin states for the genomic regions analysed samples. Robust unsupervised k-means clustering (23) is performed iteratively on this Chromatin Sample Space for a range of pre-specified number of groups (from 2 to 50). Finally, optimal clustering solutions are detected as those maximizing the Calinsky’s and Harabsz’s (CH) index (24). In an analysis involving samples from different healthy cell types, as the one presented in this work, this protocol is intended to recover those cell types, or groups of cell types, whose epigenomic differences are able to discriminate them. These epigenetically robust groups of samples allow us to confidently address the detection of those regions that are important for establishing segregation of these samples.

A challenge of this approach was to deal with millions of regions within the same analysis. However, many of these regions will not be informative for discriminating the sample groups in our dataset. Highly variable regions and completely conserved regions are non-informative regions that increase the computational time cost, while sample-specific divergent regions can bias the results, being strongly influenced by the presence of sample outliers or sample-specific experimental noise. In order to reduce the influence of sample-specific patterns contributing to outlier effects, we focused on the set of regions presenting at least two different chromatin states in at least two samples each of them. Additionally, we filter out all the regions with change patterns (vectors of chromatin states for each genomic region across samples) that were poorly represented in our dataset. In particular, we filtered out those regions whose patterns were not shared by 10 regions (we obtain similar results for patterns shared by 5, 10 and 15 regions; data not shown). This step dramatically reduces the computational burden by removing regions with low influence in sample clustering. As a result of this filtering we run our MCA framework with 275,825 regions from the 22 autosomal chromosomes of all the healthy samples.

### Selection of Chromatin Determinant Regions in the Chromatin Region Space

Concomitantly to the detection of sample clusters, ideally equivalent to cell types, our framework allows the detection of the subset of regions better reflecting this inter-sample clustering. We called these regions Chromatin Determinant Regions and they are methodologically equivalent to the Specificity Determining Positions detected (SDPs; (21)) in protein families. First, we project the vectors reflecting every genomic region/state combination into the MCA space, generating the Chromatin Region Space. Vectors representing chromatin patterns perfectly associated to every combination of sample clusters were used as fingerprints of the corresponding grouping. Every epigenomic region was associated to the closest fingerprint in the Chromatin Region Space. Finally, CDRs were defined as those positions for which all their chromatin states were among the top 10 shortest distances to its fingerprints and the combination of these fingerprints form a perfect partitioning of the sample clusters (for a more detailed description see (21)). In this situation, CDRs correspond to those regions with patterns of chromatin states along samples with very few intra-cluster epigenomic changes but with at least two clusters with different states. This definition of CDRs highlights the two key properties of these regions: the stability of their state is important for every single epigenomic cluster of samples and they define inter-cluster epigenomic changes. These properties point to the putative role of these regions in cell identity and cell fate respectively.

### Chromatin Determinant Regions annotation, expression and enrichment analyses

Genomic annotation was carried out with Hypergeometric Optimization of Motif EnRichment (HOMER software v4.7.2;(25)). The tool annotatePeaks.pl was used with default parameters to annotate CDRs to genes with the following priority assigned: TSS (from −1kb to +100bp), transcription termination site (from −100bp to +1kb), protein coding exon. 5′-UTR exon, 3′-UTR exon, intro and intergenic. More detailed information is available in http://homer.salk.edu/homer/ngs/annotation.html. Gene Ontology (Biological Process;(26)) and Reactome (27) enrichment analysis were done adding the -go flag to the annotatePeaks.pl tool. Then, we calculated a p-adjusted value based on Benjamini-Hochberg correction using R. All terms with an adjusted p-value < 0.05 were considered significant. We summarized the Gene Ontology (Biological Process) significant terms with REVIGO (28).

The expression associated analyses were carried out retrieving the RNAseq data for 60,483 protein-coding, ncRNA, pseudo, snoRNA and snRNA genes from The BLUEPRINT Data Analysis Portal (29). We took information from 12 macrophages, 8 monocytes, 6 neutrophils, 4 naive B cells, 3 germinal center B cells and 21 T cells, no data was found for HSCs from mature samples. We applied an ANOVA test to 7764 genes with CDRs associated and adjusted the p-value with Benjamini-Hochberg correction, p-value<=0.05 was considered significant. Statistical analyses were carried out with aov and p.adjust functions from R (v3.2.2).

The transcription factor motif enrichments were performed with the findMotifsGenome.pl tool included in HOMER software (v4.7.2; (25)). To determine the relative enrichment of known TFMs we excluded the CDRs referred to transcription, as they are related to polymerase elongation and not to transcription factors binding. The searches were done against a selected random background of windows adjusted to have equal GC content distribution in each of the input sequences. The region size was set up to “given”, other parameters were used by default. More detailed information is available in http://homer.salk.edu/homer/ngs/peakMotifs.html. The transcription factor motifs with a q-value<0.01 at least in one cell type were considered significant and selected to generate Fig. 3C. We did not find enriched TFMs for T-cells and neutrophils. The expression analyses for 28 of the transcription factor binding proteins of Fig. 3C were performed with the same approach described above, the transcription factors in HSCs were not included in the expression analysis since BDAP doesn’t provide data for HSCs.

Chromatin state transitions among cell types were represented with a Sankey diagram in Fig. 3A using the “makeRiver” and “riverplot” functions included in the “riverplot” R package. (v0.5; https://cran.r-project.org/web/packages/riverplot/index.html)

### Chromatin Determinant Regions in the context of disease

The “healthy” hematopoietic chromatin sample space provides us a reference sample space, reflecting the informative epigenomic distances between normal hematopoietic cell types. As it is based on the major sources of information involved in hematopoiesis, it also serves us to study to what extend leukemic epigenomes retain features important to define the cell identity of the normal cell types.

In order to get a clearer view of these residual signals of “normality”, we focused on those CDRs for which the tumoral sample shows a chromatin state present in any cell type. For this, we projected the leukemic samples on the “healthy” hematopoietic chromatin sample space, but considering only the influence of these CDRs. In practice, it means that every leukemic sample is projected based on a different number of regions and its position reflects the extent to which these regions correspond to patterns more related to one or other healthy cell type. This approach allows us to reduce the effect of tumor-related epigenetic changes and to weigh the contribution of patterns of chromatin states associated to more than one cell type according to their influence in the “normal” chromatin sample space. We also projected the prototypic “normal” cell types represented by the vectors presenting the chromatin states characteristic of the corresponding cell type for each CDR. Distances of leukemic samples to these prototypic “normal” cell types reflect the similarity of the chromatin states in CDRs balancing the effect of chromatin states shared with other “normal” cell types.

Despite the effect of focusing on “conserved” states in CDRs, highly transformed leukemic samples could include a relevant number of changes to chromatin states characteristic of a different cell type. These effects will contribute to leukemic samples with less “cell type-specific” CDRs. This situation can lead to less well-defined clusters of leukemic samples. Therefore, we decided to perform a hierarchical clustering (using Ward’s method with euclidean distances as implemented in pheatmap v1.0.8 R package, http://CRAN.R-project.org/package=pheatmap) in this CDRs-based chromatin sample space, to illustrate the association of different leukemias to different cell types. As HSCs, Macrophages and gc B cells “prototypic” cell types were clearly very distant to the projections of all leukemic samples in this space, they were not considered in the hierarchical clustering, in order to improve the resolution of the relationship of tumoral and healthy samples.

We also define the ratio of CDRs with a chromatin state different to any healthy cell type as the tumoral epigenomic divergence. It represents how divergent a tumoral sample is from the space of healthy states calculated with the normal samples. Therefore, higher divergences imply higher probabilities that the cell type of origin of the tumoral sample is not represented in the healthy chromatin space or that the tumoral sample diverged so much than its projection on this space should be taken with care. The analyzed tumoral samples show epigenomic tumoral divergences ranging from 0.02 to 0.08 with higher values for AML samples, suggesting that they can be confidently analysed in this space.

We defined CDRs altered in leukemias as those CDRs in which more than 50% of the tumoral samples show a chromatin state not observed in any normal sample. One of the advantages of this definition is that it is agnostic about the cell of origin of the tumor. Obviously, this definition, as any other, is limited to the cell types included in the study and some of these regions could be reclassified when more cell types (especially progenitor cell types) are available. In absence of more information, this criterium provides a simple definition of regions that are potentially important for tumoral progression.

Specifically altered regions in AML (or CLL or MCL) were defined as those CDRs with more than 50% of the AML (or CLL or MCL) samples presenting an unobserved state in normal cell types, but lower than 50% in the other two leukemia types. In both cases, CDRs altered in leukemia were analysed using HOMER, as explained above. For exploring tumor-specific GO and Reactome enrichments, those terms enriched also in the whole set of CDRs were filtered out from altered CDR enrichments.

### Resource

**Method availability**: https://github.com/david-juan/ChromDet

**UCSC track hub** to browse the CDRs and the chromatin states for all samples: http://genome.ucsc.edu/cgi-bin/hgTracks?db=hg19&hubUrl=http://mcahematopoiesis.bioinfo.cnio.es/carrillo_et_al_NAR/hub.txt

## Results

### The chromatin space of human hematopoietic differentiation

We carried out a multi-group comparative analysis of chromatin states for representative cell types of the myeloid and lymphoid lineages to understand how epigenetic changes in chromatin are related to hematopoietic differentiation in humans. We focused our analysis on a set of 42 blood IHEC epigenomes from eight different cell types, with at least three independent biological replicates available: hematopoietic stem cells (HSC; n=3), neutrophils (n=5), monocytes (n=6), macrophages (n=4), naive and germinal center (GC) B-cells (n=4 and n=3) and CD4 and CD8 T-cells (n=13 and n=4), see Fig. 2A and **S1 Table** for details.

These epigenomes were assembled from ChIP-seq data generated by three IHEC consortia: BLUEPRINT (n=22), NIH ROADMAP (n=18) and ENCODE (n=2). We integrated ChIP-Seq data experiments for the six core histone modification marks that are required to be included in IHEC epigenomes: H3K27ac marking active regulatory regions, H3K4me3 marking promoters (30, 31); H3K4me1, related to enhancers (30); H3K36me3, marking transcription (30); H3K27me3 and H3K9me3, associated with polycomb and heterochromatin repression, respectively (30). Importantly, we only used histone mark sets where all six marks were profiled in the same individual (i.e. each epigenome corresponds to a unique individual).

A multivariate Hidden Markov Model (HMM) was employed to learn combinatorial chromatin states based on the six histone marks using ChromHmm (18). However, others methods to segment the genome based on histone marks (or other features) could be used at this step, like Segway (32), EpicSeq (33), hiHMM (34), chromstaR (35), IDEAS (36) and others. In fact, the input for our method are the genome segmentations for the included samples. This means that users could use our method with the genome segmentations obtained by the software of his/her choice.

Further, the genome of each sample was segmented using the 11 combinatorial chromatin states model generated (see **S1 Fig**). To facilitate biological interpretation, the 11 chromatin states were further collapsed into 5 functional chromatin states encompassing five main categories: transcription (T), heterochromatin/low signal (H), repressed promoter (R), enhancer (E) and active promoter (A; see Methods for details; **S1A Fig**). Thus, for each sample, we can create a vector representing the chromatin state of consecutive 200 bp windows along the whole genome, using this reduced 5-state alphabet. In order to reduce biases associated to the different size of each regulatory region, we collapse contiguous 200 bp windows having the same chromatin states pattern along all the samples (see Methods for details).

Our initial aim was to generate a low-dimensional chromatin space, a graphical representation of the structure and dimensionality of a complex and large data set, reflecting the major sources of epigenetic differences among hematopoietic samples (eg. changes in chromatin states). To this end, we applied a protocol based on Multiple Correspondence Analysis (MCA), which we have previously applied to protein sequence (21) and gene expression (22) analysis. MCA is an analysis similar to Principal Component Analysis (PCA) but appropriate for categorical data. We created an MCA-based multi-dimensional space in which the different samples are placed based on their vectors of chromatin states across the genome. The first stage of our protocol selects the minimal number of the most informative components that are relevant in this space, which already allows us to detect clusters of samples (see Fig. 1).

**Fig. 1.**
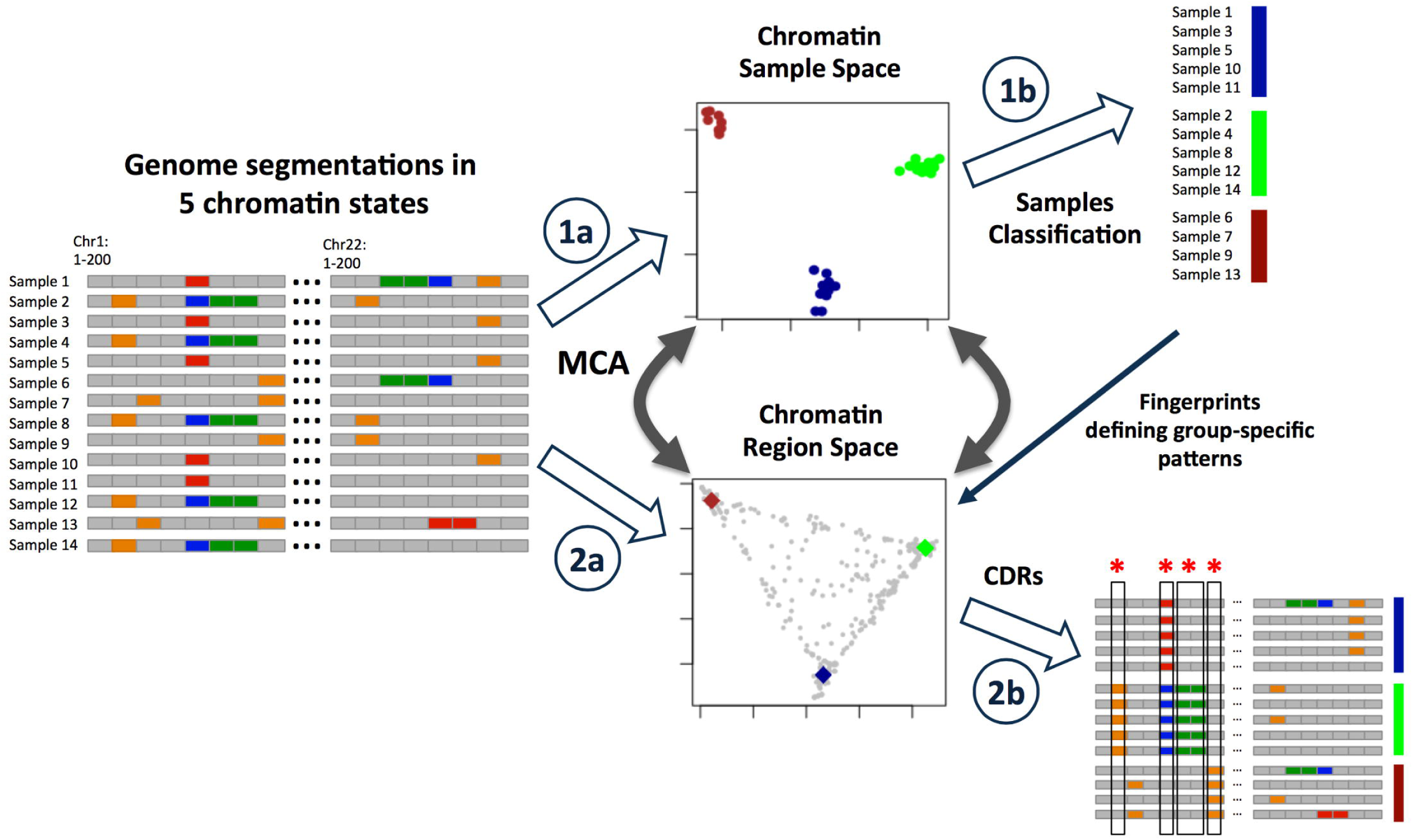
Framework to identify CDRs that determine cell or lineage identity based on chromatin state changes. 1A) A chromatin samples space is generated with MCA from the chromatin segmentations by each sample. 1B) Samples are classified depending on clusters derived from the MCA analysis. 2A) A second space is generated with MCA from the chromatin segmentations by each sample. 2B) The CDRs are obtained selecting those genomic regions that overlap with the cluster sample fingerprints, a reference sample representing each cell type cluster. These regions discriminate the different cell types classified in the samples space. (*Regions with chromatin changes among cell types -> CDRs) See also S1 Fig and supporting information

**Table 1.**
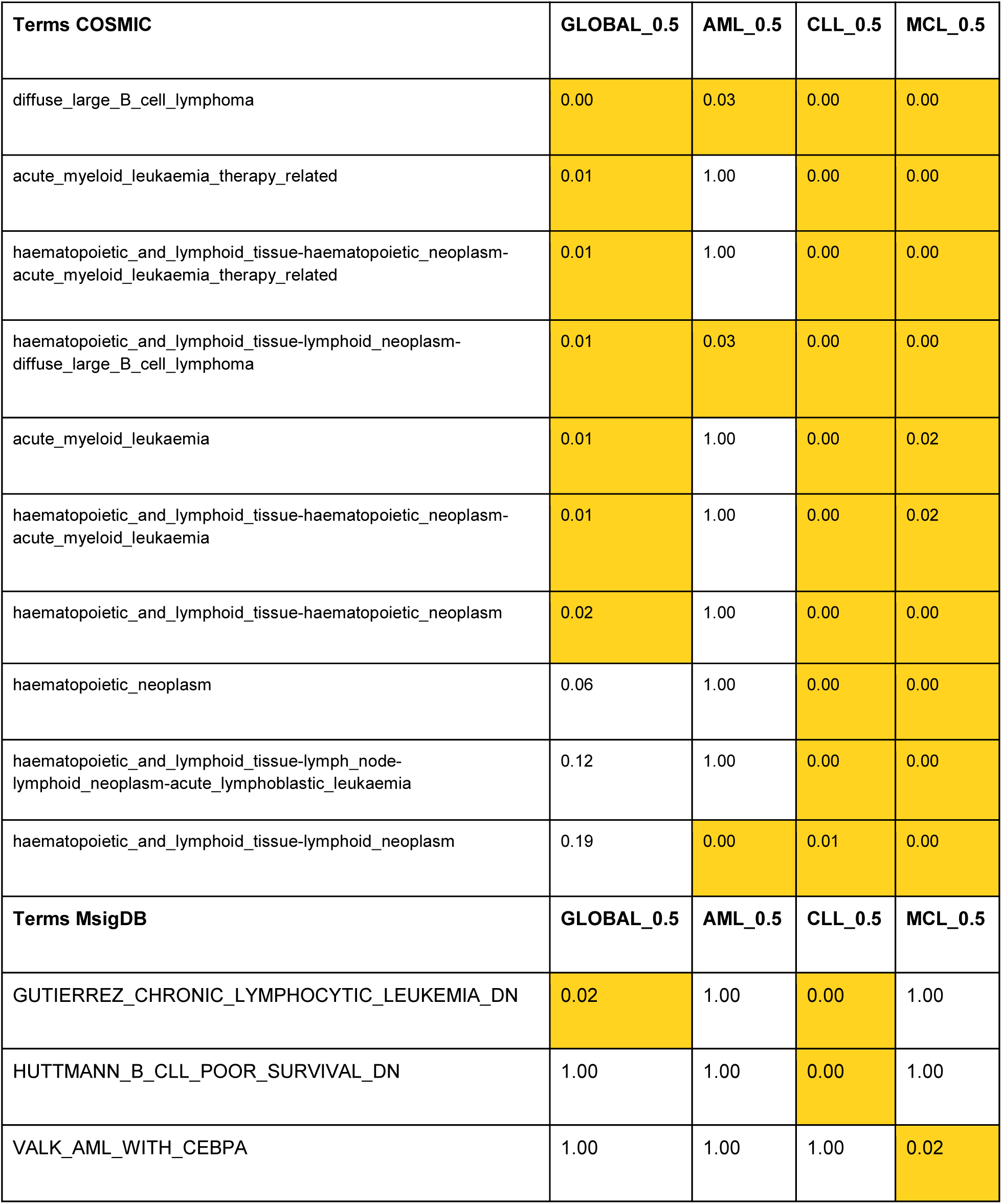
COSMIC and MSigDB enrichments for leukemia-altered CDRs. In orange background those statistically significant. See also S10 Table

Application of this approach to the matrix of collapsed chromatin states along the autosomal chromosomes in the 42 different samples results in a *hematological chromatin sample space* with the first two components as significantly informative according with a Wilcoxon test (Fig. 2B; see Methods for details). Samples from the same cell type cluster together and the major blood cell types are clearly separated from each other, showing that the origin and technical biases of the samples are not affecting the results (3 different consortia and therefore different laboratories). The relative samples distribution and the clustering are robust, as shown by analysing each of the autosomal chromosomes independently (see **S2 Fig**).

**Fig. 2.**
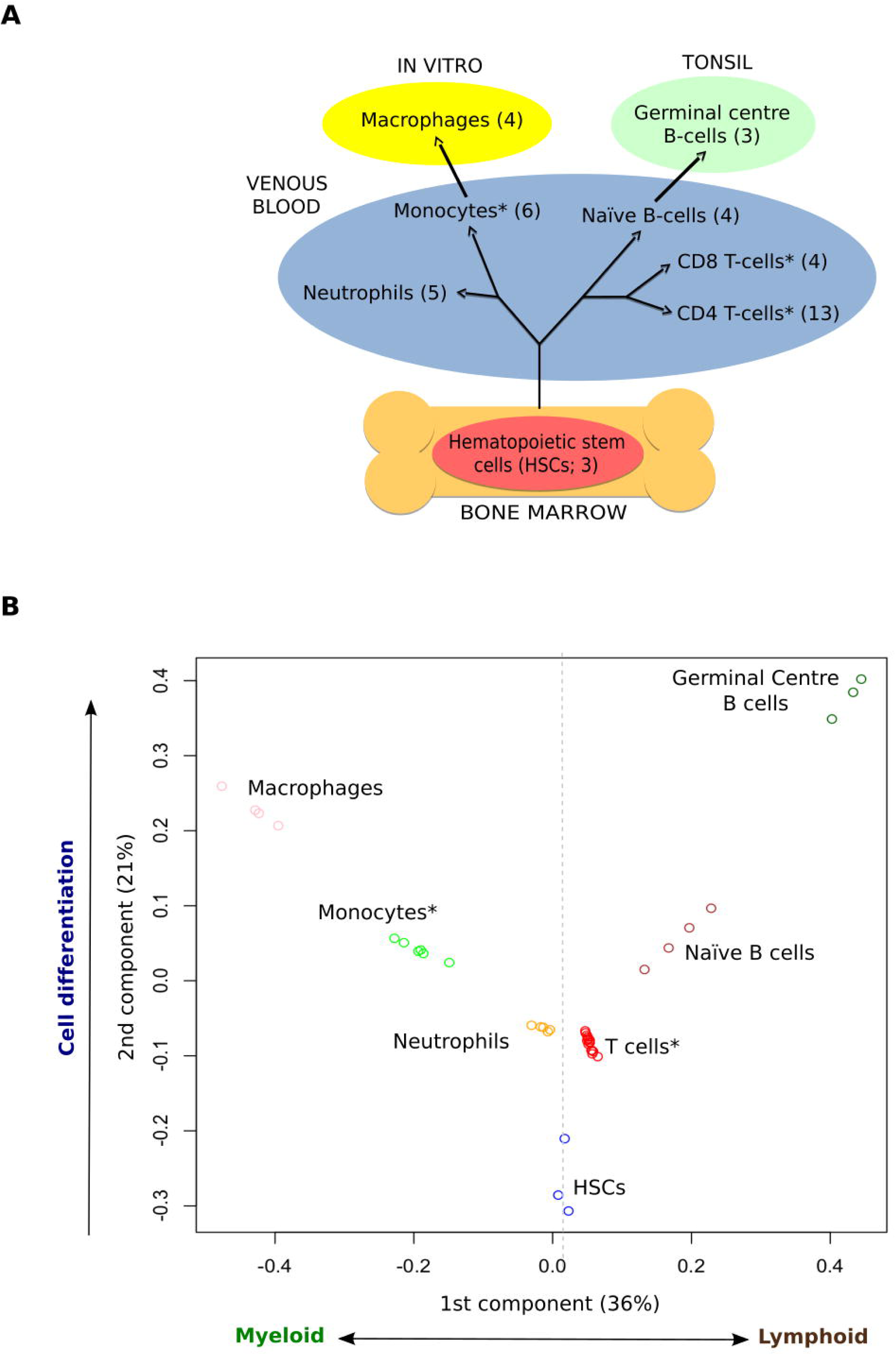
Hematopoietic cell types cluster based on chromatin states. A. Schematic differentiation tree of the cell types considered, highlighting the tissue of origin and environment of each sample type. B. Clustering of the samples in the MCA space recovers ontological relationships among cell types. (*Cell types with samples from different consortia) See also S2 Fig, S1 Table and supporting information

As in PCA approaches, the interpretation of the two components selected by our method to separate the different cell types can lead to biological insight. Interestingly, the first component, represented on the horizontal axis, clearly separates myeloid (left side) from lymphoid cell types (right side) with HSCs situated in a central position. On the other hand, the second component on the vertical axis seems to reflect the lineage-independent epigenomic changes needed for the differentiation of the cell types from the HSCs, combined with the sample environment. We can draw a path from the pluripotent HSCs in bone marrow (at the bottom of the plot) all the way to the more mature cell types or subpopulations, such as *in vitro* cultured macrophages and germinal center B cells from tonsil (at the top of the plot). The central location of neutrophils, monocytes, T cells and naïve B cells from venous blood in this space suggests less epigenomic changes between these cell types and the HSCs (see Fig. 2B). Interestingly, neutrophils and T cells are the cell types with least epigenomic changes from the HSCs. However, as in previous works based on single chromatin marks (37), we fail to discriminate CD4 and CD8 T-cells, which form a tight cluster. In conclusion, our approach is able to capture the main biological differences between cell types, and is fully consistent with the known underlying biological process, showing that epigenomic states are an excellent source of information for discriminating these cell types.

Obviously, the value of the results obtained by our approach depends on the input information. Therefore, we strongly encourage introducing proper quality criteria to decide the inclusion of a sample in the analysis (see **Material and Methods**). Interestingly, a detailed evaluation about the effect of using different input data from the segmentations (see **supporting information** and **S3-S5 Figs**) supports the use of collapsed chromatin states to discriminate samples by cell type. These analyses identify elongation and enhancer states as the most informative sources of information, and illustrates the potential of our MCA-based approach for dealing with epigenomic data. Consequently, for studying CDRs associated to differentiation, we strongly recommend collapsing chromatin states into a small number of robustly defined states reflecting major functional shifts in transcription and enhancer activities, instead of more dynamic variations in the strength of the signal associated to these functions.

### Chromatin determinant regions (CDRs)

So far we have shown how the MCA approach permits the generation of a space in which to robustly locate the different hematopoietic samples. Next, we aimed to identify the specific genomic regions that contribute most in defining specific cell types. We call these regions Chromatin Determinant Regions (CDRs; Fig. 1).

In order to retrieve these CDRs we applied the second stage of our MCA-based protocol (21). This involves building a *hematological chromatin regions space*, in which each genomic region can be located based on its patterns of chromatin states across cell types (see Fig. 1). For this we projected the chromatin states of every region of the genome on the same principal components of the Hematological Chromatin Samples Space. In this space we identify which regions have chromatin states that can discriminate the different cell types classified in the samples space (that is the different sample clusters). In practical terms, using this approach we find the CDRs that give rise to differences between cell types. For instance, a given region can show an enhancer state in lymphoid cell samples and a heterochromatin/low signal state in the rest of the samples. In other cases, our protocol allows us to recover more complex patterns, such as those in regions able to discriminate more than two cell type groups. Starting from a total of 2,687,482 genomic regions for the 22 autosomal chromosomes included in the analysis, we recovered a total of 32,662 CDRs comprising 20,421,600 bp (a 0.71% of the canonical autosomal chromosome size) (see **S4 Table**).

As mentioned above, each CDR can be associated to a pattern of states across the different cell types, pointing to chromatin changes that might be drivers of cell differentiation. The most abundant CDR patterns we identified correspond to regions that have a transcription or enhancer state in one or two cell types, while having a heterochromatin/low signal state in the others (see **S6 Fig** and **S5 Table**). The six most frequent patterns, that together comprise 61% of the CDRs, present transcription or enhancer states in GC B cells, HSC and macrophages, while having heterochromatin/low signal states in all other cell types (see **S6 Fig** and **S5 Table**). In general, CDRs related to Transcription states are larger than the ones showing patterns with other states (see **S7 Fig**). In addition, we can distinguish patterns that are cell type-specific (69,3%), lineage-specific (16,9%), which are shared by two or more close cell types, and others with more complex patterns between more distant cell types (13,8%; see **S8 Fig**). We have included UCCS browser’s screenshots of two interesting genes that show nearby CDRs, ABHD16B that shows transcription in the lymphoid lineage and LINC00494 active only in B cells (**S9 Fig**). As far as we know, these genes have not been previously related with hematopoiesis.

Recently, Corces et al (38) generated ATAC-seq profiles to analyse the chromatin accessibility in a comprehensive collection of hematopoietic cell types, of which HSCs, B-cells, T-cells and monocytes are also included in our analysis. Around 10% of their defined set of 774 cell-type-specific regions based on differential accessibility (38) overlapped with our defined CDRs. For these regions, we analysed the ATAC-seq signal distributions for different CDR patterns (see **S10 Fig**). Importantly, we found that CDRs that show cell type-specific active state patterns in HSCs, B-cells, T-cells and monocytes respectively also show increased chromatin accessibility specifically for those cell types in the ATAC-seq data.

### CDR chromatin state transitions across hematopoiesis

A more detailed analysis of the CDR transitions between cell types following the differentiation process can provide insights about chromatin remodeling across lineages. From the first pluripotent stage (HSCs), 4 possible second stages can be obtained (Monocytes, Neutrophils or Naive B cells, T cells, according to the branch). After a further round of differentiation the third stage comprises Macrophages (originating from Monocytes) and GC B cells (originating from Naive B cells). Fig. 3A shows transitions in CDR states across the various branches of the differentiation process. We observe the transitions from the HSCs to the second stage to be characterized by a turning off of active and enhancer CDRs. In contrast, in the second round of differentiation (from Monocytes and Naive B cells to Macrophages and GC B cells, respectively) there is an increase in the activation of promoter and enhancer CDRs.

**Fig. 3.**
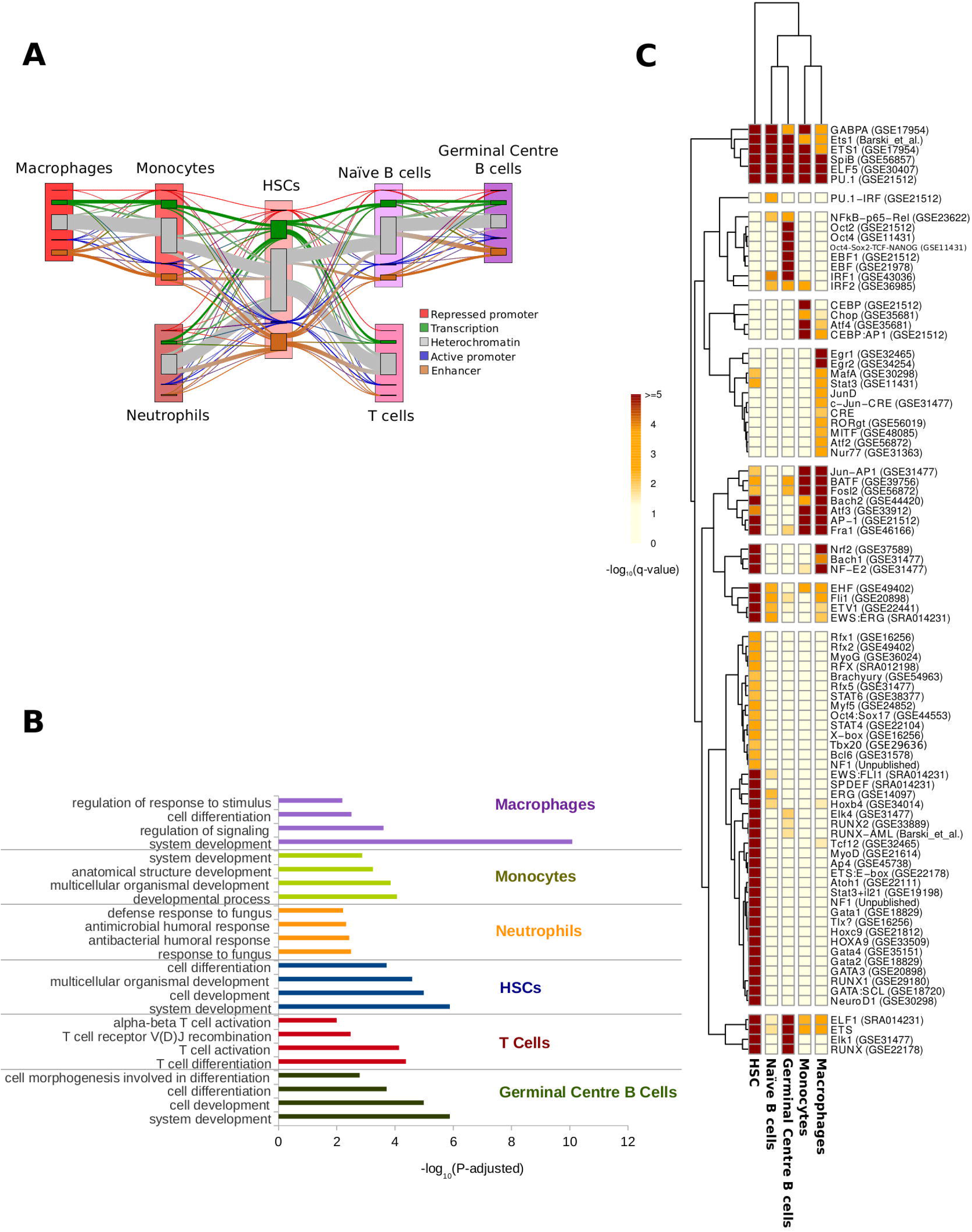
Functional and Transcription Factor binding motifs characterization of Chromatin Determinant Regions. A) Sankey Plot representation of chromatin state transitions at CDRs during hematopoietic cell differentiation. Nodes for each cell type represent the five “collapsed” chromatin states (see Methods). For each pair of cell types in the hematopoietic differentiation pathway, flows, represented by line thickness, are proportional to the number of regions that show a transition between a particular pair of states. Changes in chromatin states between two stages of differentiation are shown with lines that change colour. The thickness of the lines is proportional to the number of regions that show a transition between a particular pair of states. B) Enriched ontology terms from the genes related to the CDRs that characterize each cell type. C) Heatmap and Hierarchical clustering based on transcription binding proteins enriched in the CDRs that characterize each cell type (see Methods). See also S3-S15 Figs, S2-S6 Tables and S1 File

### CDR association to genes and transcription factors binding sites

Chromatin state changes at CDRs might be pointing to drivers of cell differentiation and could be involved in regulating the expression of nearby genes that are important for these cell type transitions. We found most of the CDRs (94,5%) in intergenic and intronic regulatory regions, with an enrichment in the promoter and 5-UTR regions over the genomic background (see **S11 Fig**). A detailed annotation of each CDR is available in **S6 Table**. We associated each cell type-specific CDR to its most proximal gene and carried out a multi-group gene expression analysis of all the mature cell types, taking advantage of The BLUEPRINT Data Analysis Portal (BDAP;(29)). The analysis was carried out on 7764 genes with gene expression data available and associated CDRs, out of 60,483 included in BDAP, including protein-coding, ncRNA, pseudo, snoRNA and snRNA genes. The analysis showed that 81% had significant gene expression differences (P-value adjusted) across the mature cell types (see **S7 Table**).

Further, functional enrichment analyses were performed for the genes associated to each cell type-specific CDR having specifically active promoter, enhancer or transcription states (see Fig. 3B; **S12-18 Figs; S8 Table**; see Methods for details). As expected, genes proximal to the CDRs defining HSCs were mainly enriched in processes related to development and cell differentiation.

CDRs defining the myeloid lineage were close to genes related to tissue development and antimicrobial response among others. On the other hand, for CDRs defining the lymphoid lineage we found genes related to T cell activation, cytokine production or response to interleukin-4, a cytokine produced by T cells involved in humoral and adaptive immunity (39). CDRs defining the two different B cell types were associated to genes with functions in proliferation and differentiation.

In addition, different neuron terms for differentiation and development were enriched for different cell types. These enrichments could be explained by the overlap in the molecular programs for hematopoiesis and neuropoiesis (40–42). The hematopoietic system is involved in many processes and genes related with neuronal development and function have been observed as expressed in different hematopoietic cell types (43). For example, we find a CDR overlapping with the gene encoding for Basp1 (Brain Abundant Membrane Attached Signal Protein) that belongs to many differentiation/morphogenesis-related GO terms, including “central nervous system development”. This and other related neuronal genes were shown to be up-regulated in germinal centre B-cells, where its pattern of gene expression is associated to the development of neurite-like projections of the membrane (44). Furthermore, interactions between the nervous and immune systems are required for organ function and homeostasis (45). A report has shown that primary CD34+ hHSCs express mRNA for a number of proteins that are used by neurons (among other cell types), including receptors for trophic factors and other mediators that are known to influence neuronal development (42). Finally, the similarity between these two differentiation programs could explain the fact that HSCs can differentiate to neural cells, albeit at relatively low efficiency (46–48).

We next asked whether CDRs involving cell type-specific active promoter or enhancer states were enriched in transcription factor motifs (TFMs, see Methods for details). Hierarchical clustering based on the TFM enrichment patterns clearly separates the HSC TFMs profiles from those of the myeloid and the lymphoid cell types (Fig. 3C). A detailed annotation for motifs in each CDR is available in **S6 Table**.

We observed in HSCs a specific motif enrichment for GATA factors, which have been related to regulation of the self-renewal of long-term hematopoietic stem cells and differentiation of bone marrow-derived mesenchymal stem cells (49–52). Enrichment in binding motifs for factors like RUNX, implied in stem cell fate maintenance and normal function, was also observed in HSCs-specific CDRs (53, 54). GATA and RUNX factors were described by Corces et al. (38) as dominant regulators of chromatin accessibility in hematopoiesis. Interestingly, motifs for the so far uncharacterized factor X gene family, known to regulate the major histocompatibility complex (MHC) class II (55), were also exclusively enriched in CDRs specific for HSCs.

In myeloid cell types, CDRs specific to monocytes are enriched in binding motifs for the C/EBP homologous protein (CEBP/CHOP) and its interactor ATF4 (56), (57), which plays a key role during the differentiation of the monocyte lineage (58, 59). In contrast, EGR1 and EGR2 binding motifs, which are essential for macrophage but not for granulocyte differentiation (60, 61), are enriched in macrophages. Higher expression at RNA level is observed in macrophages compared with monocytes and mature neutrophils (see **S19 Fig**). In addition, enrichment for transcription factor binding sites related to macrophage differentiation like STAT3, JUND, MITF, NUR77 or ATF2 is observed in CDRs specific to macrophages (62–66).

Binding motifs for members of the NF-KB complex (NF-KB, RELA, IRF2), implicated in stimulus response, were enriched in CDRs characterizing GC B cells. It is known that defects of this complex in germinal centers affect their maintenance and B cell differentiation (67, 68). In addition, we observed enriched motifs for Early B-cell factor 1 (EBF1), a central transcription factor in B cells implicated in germinal center formation and class switch recombination (69, 70), Oct2 and Fli1, transcription factors expressed in B cells and related to normal B cells proliferation (71, 72).

TFMs from the ETS transcription factor family genes (GABPA, ETS1, SpiB, PU.1 and ELF5) were enriched in all cell type-specific CDRs. These gene families are ubiquitously expressed in the different blood cell types, although they are known to play specific roles in different cell types. For example, in monocytes, PU.1 regulates the transcription of a large proportion of myeloid-specific genes, while in B cells it is involved in regulating the transcription of the heavy and light immunoglobulin chain genes (73).

Finally, we took advantage of BDAP expression data for 28 transcription factors whose DNA binding motifs were enriched in CDRs (Figure 3C) and for which expression data was available. We excluded transcription factors whose binding motifs were specifically enriched in HSCs as this immature cell type is not included in BDAP. A subset of 96,5% (27/28) of them showed differential expression between cell types (see **S7 Table**). In addition, we observed that 60% (16/27) of the transcription factors with changes in expression also have a CDR associated to them by proximity, suggesting a central role for chromatin regulatory regions in the hematopoietic regulatory network.

Taken together, the gene expression, gene ontology and TFM enrichment analyses suggest that the identified CDRs are indeed important functional regions, where chromatin remodeling is linked to cell fate. Overall, we have shown that our approach is useful to identify key and potentially driver local changes in the epigenomes of healthy cells across different hematopoietic lineages.

### Clustering of healthy and leukemic samples based on CDRs

The framework explained above allowed us to identify specific genomic regions that are under epigenetic control and might contribute to define blood cell types. This framework can be further exploited to analyze the relationships between leukemia and healthy cell types.

Extensive epigenetic changes are common in most leukemias and solid tumors (74) and epigenetic features such as DNA methylation or open chromatin have been shown to be useful to identify the cell of origin of tumours (75, 76). However, given the extensive genome-wide epigenetic alterations of tumour cells, matching tumoral cells with their healthy counterparts is a great challenge and an essential step to identify the chromatin changes leading to malignancy.

The CDRs constitute an epigenetic signature of hematopoiesis. Therefore, we reasoned that they should be useful to classify blood cancer samples according to their similarity to normal cell types. We used the data generated by The BLUEPRINT consortium for three hematopoietic neoplasms, including 6 chronic lymphocytic leukemias (CLLs), 3 acute myeloid leukemias (AMLs) and 3 mantle cell lymphomas (MCLs) to explore the epigenetic similarity among healthy and cancerous samples.

We projected the leukemic samples on the healthy hematopoietic chromatin space, based on their chromatin states at CDRs (see **S20 Fig** and Methods). Next, we used the distance of each leukemic sample to a reference healthy cell type (**S20 Fig**) to quantify the similarities and differences observed at the CDRs level between healthy and disease epigenomes.

The distribution of the leukemia samples in the CDRs healthy hematopoietic chromatin sample space separates them into two main groups. The AML samples localized into the myeloid region of the space, while the CLL and MCL samples were in the lymphoid region (see **S20 Fig**). A hierarchical clustering based on the distances of each leukemia sample to each reference healthy cell type shows that CLL and MCL samples both cluster with the reference Naïve B cell (see Fig. 4; cluster I). In contrast, AML samples are distributed in more than one cluster, with two samples clustering within the reference neutrophil cluster IV, and the other one within the reference monocyte cluster II, suggesting a different origin for these tumours.

**Fig. 4.**
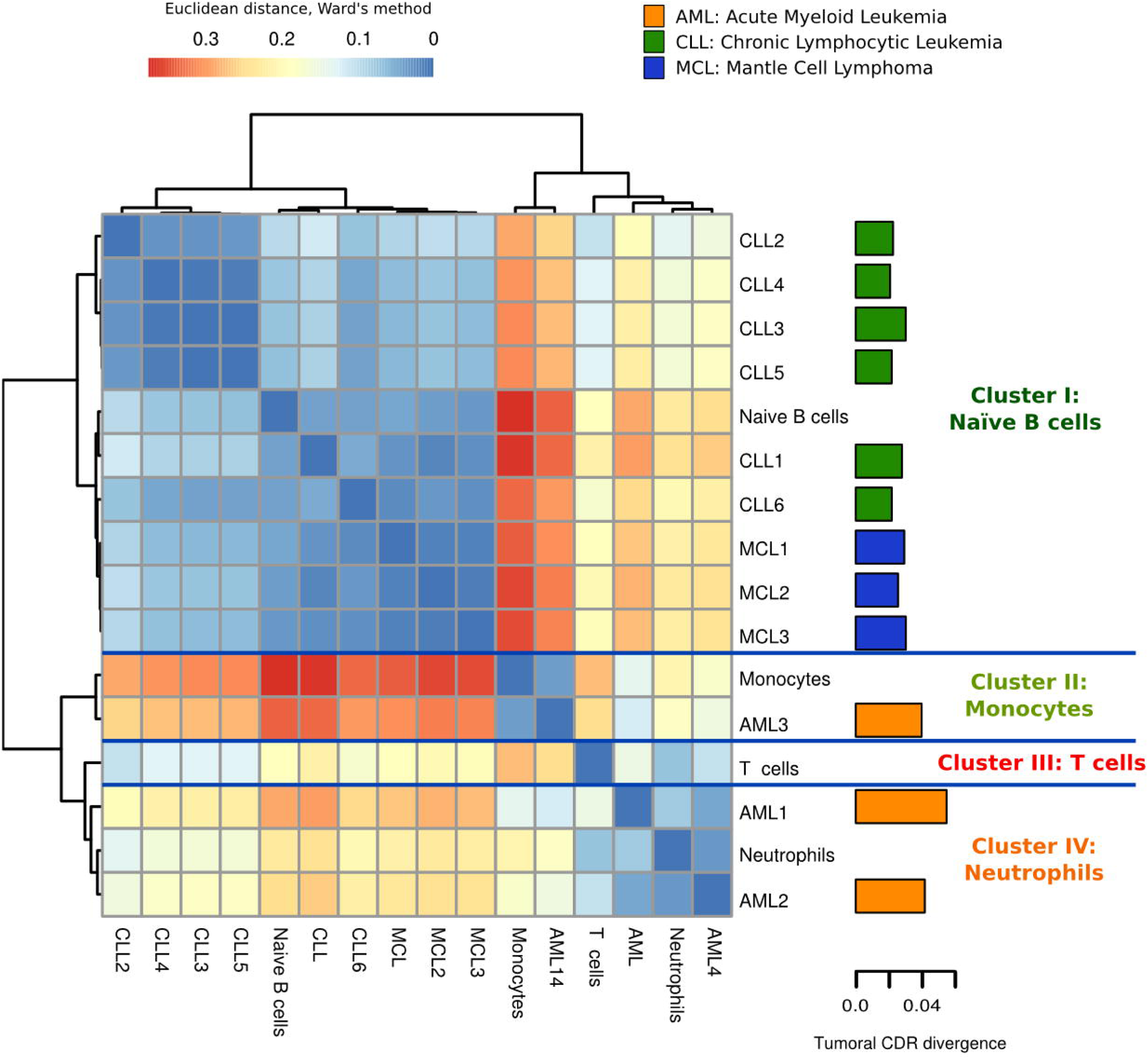
Hierarchical Clustering of leukemias based on CDRs of healthy cell types suggests potential lineage origin of tumors. The healthy cell type clusters are summarized by each fingerprint, a reference sample representing each cell type cluster. The euclidian distances between samples and fingerprints are calculated with the Ward’s method. The barplot in the right shows the epigenomic divergence (ratio of chromatin changes in CDRs) of each cancer sample to the healthy states. See also S16-20 Figs and S7-S9 Tables

Each tumoral sample was projected onto the healthy hematopoietic chromatin sample space using the CDRs whose chromatin states are represented in any of the healthy cell types (see Methods). However, there is a variable number of CDRs per tumoral sample whose chromatin state is not represented in the normal cell types. We can view these chromatin states either as features related to maturation stages of cells not included in our analyses, or as changes that have occurred specifically in the malignant transformation. Interestingly, we can observe characteristic divergence patterns for the different neoplasms (Fig. 4). AML samples appear to be epigenetically more divergent from the healthy states than those closer to the B cell derived cancer samples.

We analysed these divergent CDRs as a potential source of information about epigenomic alterations that might be important for tumoral transformation. To this end, we focused on those 297 CDRs where most of the leukemic samples have a potentially unhealthy chromatin state (a chromatin state that is never observed in the healthy samples, see **Methods and S9 Table**). We proceeded by associating these CDRs to genes by proximity and investigated whether these genes were commonly regulated or mutated in tumours.

Functional enrichment analysis of the 177 genes associated to these CDRs by proximity showed that they are mutated in a large number of tumors, including AML and CLL ( COSMIC tumoral signatures (77), p-value<0.05, see **S10 Table**). An analysis of those CDRs specifically altered in each of the leukemias (282 CDRs in AML, 591 in CLL and 727 in MCL) shows similar results. However, we observed that mutational signatures associated specifically to AML, CLL and MCL were enriched only in tumors different from the leukemia where we detected the epigenomic alteration (see **S10 Table**).

On the contrary, when comparing with gene signatures regulated in tumours, we found that genes associated to altered CDRs in AML and CLL respectively are enriched in the expression signatures of the corresponding leukemias (MSigDB gene expression signatures (78), pvalue<0.05, see **S10 Table**). This result supports that these alterations in CDRs are linked to the detected gene expression changes in the associated genes (see **S10 Table**).

We also found that the three sets of genes specifically altered in the different leukemias are all enriched in the same general processes: differentiation and development, cell-cell adhesion, endocytosis and phagocytosis or metabolic processes (GO biological process, p value < 0.05, see **S11 Table** and **S21-24 Figs**). Although these sets of genes are related to similar processes, they contain different genes (only three genes in common among the three leukemias) and they are related to different detailed functions. In fact, genes associated to CDRs altered in most AML samples are mainly enriched in membrane transporters and metabolic pathways, those altered in CLL are enriched in many signal transduction pathways (VEGF, WNT, FGFR, ERBB or MAPK signalling) and those in MCL in morphogenetic and developmental processes (p value<0.05, see **S11 Table**). These observations draw a scenario where leukemic mutations and epigenomic alterations point to the same processes that are key for tumor progression, but involve different genes in a leukemia-specific way. Taken together, these results show the potential of our proposed CDR approach to characterise hematopoietic cell types in normal differentiation and disease.

## Discussion

Chromatin remodeling is an essential process for determining the set of phenotypes deployed by eukaryotic cells. Chromatin regulation is based on combinatorial associations among proteins and complex communication networks, which define the functional states of the different genomic/chromatin regions (12). These functional states play a determinant role to define cell identity during the differentiation process. Despite the great efforts made in the last few years to generate functional chromatin maps for many cell types (19, 37), we are still far from identifying the genomic regions where driver chromatin changes occur, their association with functional changes that give the cell its identity during development, or their implications in disease.

Hematopoiesis is possibly the best characterized differentiation process, usually represented by a hierarchical tree based on morphological criteria and refined with surface markers (1). Hematopoiesis provides a well-defined model to study cell differentiation from an epigenetic perspective. We face the challenge of studying this process by integrating epigenomic information from multiple human blood cell types and different data sources. The blood IHEC epigenomes provide a unique opportunity to investigate the epigenetic basis of lineage determination.

We have developed a new protocol, based on a useful and powerful multivariate framework based on a rigorous statistical approach, to define in an unsupervised manner which cell types are epigenetically distinguishable. Importantly, we simultaneously identify the key genomic regions driving these differences. These regions, named Chromatin Determinant Regions (CDRs), can be considered as the epigenetic signatures of human hematopoiesis, a set of reference regions that through their epigenetic changes might be able to drive hematopoiesis.

The results are robust to the possible noise introduced by consortia-specific protocols and the clusters obtained provide perfect classification of samples in the different cell types. We observed clear clusters for seven cell types plus an additional cluster for CD4+ and CD8+ T cells. Interestingly, a recent work using H3K4me1 and H3K27me3 histone modifications independently was also unable to discriminate CD4+ from CD8+ T cell types (37), supporting the hypothesis that the epigenomes of these cell types are very similar.

The sample space, in addition to clearly separating the myeloid from the lymphoid lineages, reflects the epigenetic distance of each cluster from the HSC. Although both the classical and the more recent alternative hematopoietic hierarchical differentiation models propose a similar differentiation distance for neutrophils and monocytes or T and B cells(1, 2), our space shows clearly very different epigenetic differentiation distances for neutrophils and monocytes, as well as for T and B cells. These differences suggest that cell types with shorter epigenetic distances from HSCs may reach the mature state earlier. In the case of murine fetal liver T and B cells, it is known that the T cell progenitors appear earlier than the B cells ones (79).

The classical hematopoietic model establishes that the HSCs differentiate into the common myeloid progenitor (CMP) or the common lymphoid progenitor (CLP), divided in the myeloid and the lymphoid lineages (1). However, this model is under discussion, as it has been shown by Kawamoto et al. (79) and other authors (80–83) that the T and B cell progenitors retain the potential to differentiate into myeloid cells. These results have led to the proposal of an alternative “myeloid-based” model for hematopoiesis (79), which would suggest that the two main branches are not as well separated as initially thought. Interestingly, we found that the epigenetic distance between neutrophils and T cells is very short in our model, both cell types being very close to the HSC group.

Unfortunately, although our CDRs refer to chromatin changes during all the lineage differentiation steps, data for progenitors (GMP, CMP, CLP, MPP, …) do not meet the IHEC standards and could not be used in our analysis. Therefore, we can not assign each CDR to the precise intermediate cell type in which it was originated. The future availability of complete epigenomes for more cell lineages, including intermediate progenitors, will provide additional information to assess whether the myeloid-based differentiation model proposed by Kawamoto et al. (79) is consistent with the chromatin landscape.

The strength of our protocol, beyond providing a classification of cell types, is to identify the CDRs that drive human hematopoiesis. We detected 32,662 CDRs that represent the epigenetic signature of hematopoiesis for the cell types included in the analysis. Interestingly, we observed that all the transitions starting from HSCs to other cell types were enriched in epigenetic inactivation, while the Monocytes-to-Macrophages and naive-to-GC B cells transitions are enriched in epigenetic activation. These results suggest that the differentiation process involves a first phase characterized by loss of stemness through epigenetic repression of the HSC processes, followed by activation of more specific regulatory programs that define specific differentiated cell types (84–87).

A further characterization of these CDRs showed that they are enriched in DNA binding motifs of transcription factors with a key role in hematopoiesis. These results support the idea of CDRs as driver regions whose chromatin reconfiguration is associated to cell type-specific regulatory programs. Moreover, we also observed that these regions are proximal to genes with functions in cell differentiation and cell type- or lineage-specific processes, coherent with the transitions reflected by the epigenetic pattern of the regions.

As only a subset of blood cell types was used in this analysis, these CDRs have to be seen as only a first approximation to understand human hematopoiesis from an epigenetic perspective. It is important to note that other previous models were proposed based on surface markers (88) or mice models with DNA methylation (4) and transcriptomics (2). Although the human hematopoietic differentiation model closely resembles the murine one, accumulated evidence has shown that they differ in important aspects. For example, the HSC immunophenotypes (1) or hematopoietic gene regulation programs are not fully conserved between species (89).

In addition to providing a useful epigenetic signature of hematopoiesis, we have also shown that the CDRs could provide useful information about disease related epigenetic features. We applied our method to study the epigenetic similarities between leukemias and healthy cell types by projecting the leukemia samples in the space generated with the CDRs. We hypothesized that leukemia derived from certain healthy cell types would maintain the epigenetic CDR signature of its cell of origin. Indeed, our approach recovers a coherent distribution of hematological cancers, with B-cell neoplasms clustering close to B naïve cells, and a more heterogeneous classification of the AML samples. AML is known to be a very heterogeneous disease with many different subtypes and a difficult clinical classification (90, 91), which would explain why two of the AML samples cluster close to neutrophils, and the other one with monocytes. In addition, we also performed a functional analysis of the CDRs more recurrently epigenetically changed in different leukemias, showing that they tend to target general processes (such as differentiation and development, cell-cell adhesion, endocytosis and phagocytosis). Interestingly, different genes within these pathways are either epigenetically altered or mutated in the specific leukemias, suggesting mutual exclusivity of the two types of alterations in the same genes. In summary, our proof of concept application of the epigenetic signature of hematopoiesis in the study of leukemia shows the power of our methodology. Only when more leukemia and complete progenitor epigenomes will become available will we be able to exploit the full potential of this approach.

In conclusion, our results have shown the value of our multivariate framework in investigating the differentiation processes. We propose a catalog of epigenetic signatures of human hematopoiesis, based on the CDRs that best describe the different cell types. This catalog, with further refinements by the inclusion of additional cell types and hematopoietic progenitors, could become the reference IHEC resource for human hematopoiesis studies.

## Funding

This work was supported by the European Union’s Seventh Framework Programme (FP7/2007-2013) [282510] (BLUEPRINT).

## Acknowledgments

We thank José María Fernández and Jon Sánchez from Structural Computational Biology Group at Spanish National Cancer Research Center (CNIO) for technical support. Vera Pancaldi acknowledges a CNIO-Friends fellowship.

